# Stress-induced brain responses are associated with BMI in women

**DOI:** 10.1101/2023.03.03.531040

**Authors:** Anne Kühnel, Jonas Hagenberg, Janine Knauer-Arloth, Maik Ködel, Michael Czisch, Philipp G. Sämann, BeCOME working group, Elisabeth B. Binder, Nils B. Kroemer

## Abstract

**Background:** Stress is associated with elevated risk for overweight and obesity, especially in women. Since body mass index (BMI) is correlated with increased inflammation and reduced baseline cortisol, obesity may lead to altered stress responses. However, it is not well understood whether stress-induced changes in brain function scale with BMI and if peripheral inflammation contributes to this.

**Methods:** We investigated the subjective, autonomous, endocrine, and neural stress response in a transdiagnostic sample (N=192, 120 women, M_BMI_=23.7±4.0 kg/m^2^; N=148, 89 women, with cytokines). First, we used regression models to examine effects of BMI on stress reactivity. Second, we predicted BMI based on stress-induced changes in activation and connectivity using cross-validated elastic-nets. Third, to link stress responses with inflammation, we quantified the association of BMI-related cytokines with model predictions.

**Results:** BMI was associated with higher negative affect after stress and an increased response to stress in the substantia nigra and the bilateral posterior insula (*p*_FWE_<.05). Moreover, stress-induced changes in activation of the hippocampus, dACC, and posterior insula predicted BMI in women (*p*_perm_<.001), but not in men. BMI was associated with higher baseline cortisol while cytokines were not associated with predicted BMI scores.

**Conclusions:** Stress-induced changes in the hippocampus and posterior insula predicted BMI in women, indicating that acute brain responses to stress might be more strongly related to a higher BMI in women compared to men. Altered stress-induced changes were associated with baseline cortisol but independent of cytokines, suggesting that the endocrine system and not inflammation contributes to stress-related changes in BMI.

## 1. Introduction

Stress is an everyday occurrence, but prolonged exposure to stress increases the risk for a number of negative health outcomes, including for metabolic and cardiovascular disease (McEwen, 2004). Importantly, chronic stress has been associated with a heightened risk for obesity (Jackson et al., 2017) which in turn is also associated with cardiovascular events (Holmes et al., 2014; Iliodromiti et al., 2018; Khan et al., 2018) as well as dysregulations of energy metabolism (Moller and Kaufman, 2005) and the immune system (Karczewski et al., 2018). Obesity is also related to altered functioning of the hypothalamus-pituitary axis (HPA, van der Valk et al., 2018) as indicated by, for example, reduced baseline HPA activity levels (Champaneri et al., 2013; Schorr et al., 2015) and possibly altered acute endocrine stress reactivity (Incollingo Rodriguez et al., 2015). This possible bidirectional link between stress and obesity may be of high relevance for the pathophysiology of a number of stress-related disorders.

Although chronic stress is a well-known risk factor for obesity especially in women (Chen and Qian, 2012), it has not been conclusively resolved whether there are replicable differences in the acute stress response with obesity. To date, most studies have focused on altered endocrine stress responses in obesity, and there is evidence that acute cortisol responses to stress are higher in obesity (Incollingo Rodriguez et al., 2015) especially in women (e.g., Benson et al., 2009; Epel et al., 2000; Tomiyama et al., 2011, for a review: Incollingo Rodriguez et al., 2015) but there are also inconsistent results in the literature (e.g., Herhaus and Petrowski, 2018; McInnis et al., 2014; Therrien et al., 2010). Likewise, higher body mass index (BMI) is consistently associated with a blunted stress-induced cardiovascular response (Burch and Allen, 2014; Jones et al., 2012). BMI-dependent differences in subjective responses to acute stress are less well characterized, although one potential link between stress and increased food intake is compensatory eating in response to negative emotions (Srivastava et al., 2021). Taken together, there is preliminary evidence for changes in acute stress reactivity on the endocrine and cardiovascular level in overweight and obesity, but little is known about underlying changes in neural stress responses.

In addition to changes in endocrine and cardiovascular systems, overweight and obesity are also characterized by increased inflammation. For instance, higher BMI has been associated with higher peripheral blood levels of cytokines such as interleukin 6 (IL-6), high sensitivity c-reactive protein (hsCRP), interleukin 1 receptor antagonist (IL-1RA), and tumor necrosis factor (TNF) (Benson et al., 2009; Caslin et al., 2016; Mathieu et al., 2010). Acute psychosocial stress also induces a response of the immune system, with transient increases in IL-6 (Benson et al., 2009; Marsland et al., 2017; McInnis et al., 2014; Segerstrom and Miller, 2004). Moreover, baseline levels of cytokines such as macrophage migration inhibitory factor (MIF) and IL-6 were associated with subsequent stress-induced cortisol responses (Edwards et al., 2010; Kunz-Ebrecht et al., 2003) highlighting the interdependence of the immune and endocrine system in orchestrating stress responses (Rohleder, 2019). Furthermore, inflammation induced by vaccination (Harrison et al., 2009) and in depression (Aruldass et al., 2021) as well as obesity (Park et al., 2020; Syan et al., 2021) have been linked to changes in functional connectivity in brain networks related to interoception and default mode processing that are also implicated in stress-reactivity and anxiety symptoms (Kühnel et al., 2022). Likewise, both obesity and mental stress-related disorders such as depression share genetic factors that are also associated with higher risk for inflammation (Kappelmann et al., 2021). To conclude, obesity is linked to inflammation that interacts with the endocrine system to tune acute stress responses and may mediate the effects of obesity on stress.

Despite the recent progress in delineating the link between the immune system and adaptive as well as non-adaptive responses to stress, whether obesity contributes to changes in acute stress reactivity and if this is modulated by peripheral markers of inflammation is not yet understood. Here, we investigated the modulating effects of BMI on subjective, autonomous, endocrine, and neural stress reactivity. To this end, we used stress-induced changes in brain response (i.e., activation changes). Moreover, we derived dynamic functional connectivity and activation trajectories within a putative stress network (Kühnel et al., 2022) and used a cross-validated elastic net to predict interindividual differences in BMI from these imaging features. We then evaluated whether BMI-associated increases in baseline cortisol or inflammation markers contributed to altered neural stress reactivity in obesity. By unraveling sex-specific associations of BMI with acute stress reactivity, we shed new light on the interrelation of stress and obesity in women.

## 2. Materials and Methods

### 2.1 Participants

For the analyses reported here, we used a subsample of 192 participants (120 women, M_age_=35.0 years±12.3 years) from the Biological Classification of Mental Disorders (BeCOME) study (ClinicalTrials.gov: NCT03984084, (Brückl et al., 2020)) that completed a psychosocial fMRI stress task and with available BMI data. In addition, all participants completed a standardized diagnostic interview (Wittchen et al., 1995) and n=82 (42%) fulfilled the criteria for at least one mood or anxiety disorder (ICD-10 code F3-F4, excluding specific phobias) within the last 12 months. Of those, n=7 reported present medication for their symptoms. Moreover, for 148 participants (N=89 women), levels of peripheral immune marker and cortisol concentrations were available. As previously reported, we excluded participants with missing or low-quality data for each analysis separately (N=148-192, (Kühnel et al., 2022)).

### 2.2 Experimental procedure

On the first study day, participants arrived at approximately 8 am. Before the start of the experiments, a blood sample was taken to assess baseline cortisol and cytokines. On the second study day and in the second fMRI session, the stress task (Figure S1) was included (Brückl et al., 2020). To assess the cortisol response throughout the task, four saliva samples were taken using Salivettes (Sarstedt AG & Co., Nümbrecht, Germany). Additionally, we assessed the serum cortisol response using blood samples in a subset of participants. The first sample was taken upon arrival (T1) which was followed by the placement of an intravenous catheter in a subsample (n=31, 16%) of participants. The second sample (T2) captured a potential cortisol response to the placement of the catheter and was taken approximately 20min later, directly before entering the scanner. The fMRI session started with an emotional face-matching task (∼12min), followed by a baseline resting-state measurement. Before the start of the stress task (T3), participants rated their current affective state by answering the *Befindensskalierung nach Kategorien und Eigenschaftsworten* (BSKE;(Janke, 1994); Supplementary Information, SI) via the intercom. We used a psycho-social stress paradigm that was adapted from the Montreal imaging stress task (Pruessner et al., 2008). In this task, participants have to perform arithmetic tasks under time pressure and with negative performance feedback (Brückl et al., 2020; Elbau et al., 2018; Kühnel et al., 2020) that is given after each trial and additionally verbally between task blocks. These tasks typically correspond to mild laboratory stressors with 47%-65% cortisol responders (Noack et al., 2019). It starts with 5 control task blocks (60s each) interleaved with rest blocks (40s) of the *PreStress* phase where the arithmetic problems are shown with sufficient time and without negative feedback. This was followed by the *Stress* phase in which the 5 task blocks (again 60s) are presented with time pressure and negative feedback inducing psycho-social stress. The task ends with a *PostStress* phase that is analogous to *PreStress* to assess stress recovery. The Total time of the task is about 25min. Throughout the fMRI session, we measured heart rate (HR) using photoplethysmography (SI) and in the subgroup with additional serum cortisol assessment two further samples were taken (T5 and T5). After completion of the task, participants rated their current affective state again using the BSKE. Thereafter, another saliva sample was taken (T6) and participants were moved outside of the scanner for a 30min rest period. To assess stress-induced changes in resting-state functional connectivity (Vaisvaser et al., 2013; Veer et al., 2011) this was followed by another 6-minute resting-state scan. To conclude the session, participants rated their affective state and gave a last saliva sample for cortisol assessment (T8).

### 2.2 Assessment of endocrine and cytokine concentrations

After collection, all saliva samples were centrifuged and stored at -80° C until further processing. Salivary cortisol concentrations were measured with electro-chemiluminescence-assay (ECLIA) kit (Cobas®, Roche Diagnostics GmbH, Mannheim, Germany). Samples (IDs) were randomized across different batches, but all samples from one participants were processed in the same batch. The detection limit was 1090 pg/mL. The %CV (coefficient of variation) in saliva samples with varying concentrations was between 2.5% and 6.1% for intra-assay variability and between 3.6% und 11.8% for inter-assay variability. Five participants had to be excluded due to insufficient saliva volume at T6.

For a subset of the BeCOME study (n=198) cytokine assays were measured together with samples from a second in-house study (Kopf-Beck et al., 2020). Blood was collected in Sarstedt plasma tubes at 8.15am in a fasted state and frozen at -80° C after centrifugation and aliquotating. The V-PLEX Human Biomarker 54-Plex Kit (Meso Scale Diagnostics, Rockville, USA) was used to measure immune markers in plasma. MSD plates were analyzed on the MSD MESO QucikPlex SQ 120 imager (MSD). Additionally, the following markers were measured with enzyme-linked immunosorbent assay (ELISA): high-sensitivity C-reactive protein (Tecan Group Ltd., Männedorf, Switzerland, Cat # EU59151), cortisol (Tecan Group Ltd., Männedorf, Switzerland, Cat # RE52061), interleukin 6 (Thermo Fisher Scientific, Waltham, USA, Cat # BMS213HS), interleukin 6 soluble receptor (Thermo Fisher Scientific, Waltham, USA, Cat # BMS214) and interleukin 13 (Thermo Fisher Scientific, Waltham, USA, Cat # BMS231-3). All assays were performed according to the manufacturer’s instructions. Cytokines measured with the V-PLEX Biomarker Kit with more than 16% missing values were excluded, resulting in 42 cytokines (Figure S2) for further analysis. For the markers measured with ELISA, values below the detection limit were set to zero.

Likewise, values above the detection limit to the assay were set to the upper limit. All remaining markers were quantile-normalized so values were first ranked and then mapped to the quantiles of a standard normal distribution using custom code. Next, data was batch-corrected for the biobank storage position with linear regression in R version 4.0.2. A linear model was fit for each cytokine to regress out the batch variable and resulting residuals were used for further analysis. Missing values were imputed 100 times with the R package mice 3.13.0, using age, self-reported sex, biobank storage position, Beck Depression Inventory (BDI-II; Beck et al., 1996), BMI and the study as covariates. For further analysis, the median imputation values were used, as the cytokines included for further analysis (i.e., related to BMI) had at most 2 imputed values (Figure S3).

### 2.3 fMRI data acquisition and preprocessing

We acquired MRI data using a 3T scanner (GE Discovery MR750). The stress task consisted of 755 T2*-weighted echo-planar images (EPI, interleaved acquisition TR= 2s, TE= 40ms, 64 × 64 matrix, field of view = 200 × 200 mm^2^, voxel size = 3.5 × 3.5 × 3 mm^3^). Both resting states consisted of 155 EPIs (TR= 2.5s, TE= 30ms, 96 × 96 matrix, field of view = 240 × 240 mm^2^, voxel size = 3.5 × 3.5 × 3 mm^3^) each. Preprocessing was performed in MATLAB 2018a and SPMv12 as previously reported (Kühnel et al., 2022, 2020). First, fMRI volumes were corrected for slice-timing. Then, to correct for head motion, fMRI data was realigned to the first image and six movement parameters were derived for later noise correction. For spatial normalization a high-resolution T2*-weighted image was first segmented using the unified segmentation approach (Ashburner, 2007). Derived grey and white matter segments were then used for normalization to the MNI-template by applying DARTEL (Ashburner, 2007). Last, data was spatially smoothed with a 6 x 6 x 6 mm^3^ full-width at half-maximum Gaussian kernel. To perform physiological noise correction, we used aCompCor (Behzadi et al., 2007). To this end, we extracted timeseries of the normalized but unsmoothed functional data of all voxels from white matter and cerebro-spinal fluid segments (probability maps thresholded at p> .90). We then performed PCA and used the first five components of each segment as physiological noise covariates.

### 2.4 Data analysis

#### 2.4.1 Stress response to the psycho-social stress task

##### Endocrine response

To assess the endocrine response to the stress task, we calculated the change in cortisol concentration between T2 and T6. To account for potential responses of the HPA-axis to the placement of the intravenous catheter (Kühnel et al., 2020), we included a dummy-coded nuisance regressor in all analyses that classified participants as pre-task cortisol responders when the concentration at T1 compared to baseline (T0) exceeded 2.5 nmol/l (Kühnel et al., 2020; Wust et al., 2000).

##### Autonomous response

To assess the autonomous stress response (Kühnel et al., 2020), we calculated the change in average HR during arithmetic blocks in the *Stress* or *PostStress* phase compared to *PreStress.* After preprocessing raw PPG data and performing beat detection using the Physionet Cardiovascular Signal toolbox (Vest et al., 2018), we derived the average HR of each task or rest block with the RHRV package (Rodríguez-Liñares et al., 2008) for R. This included further preprocessing to exclude implausible interbeat-intervals (IBI). We filtered out IBIs shorter than 0.3 s and longer than 2.4s and excluded IBIs showing excessive deviations from the previous, following, or running average (50 beats) IBI. The threshold for excessive deviations was updated dynamically with the initial threshold set at 13% change from IBI to IBI (Vila et al., 1997).

##### Affective response

To assess the subjective stress response, we calculated the change in positive (activity, wakefulness, self-certainty, focus, and relaxed state of mind) and negative (internal and external agitation, anxiety, sadness, anger, dysphoria, sensitivity as well as three items assessing somatic changes) sum scores form the respective items directly after the task (T6) and after the 30-minute rest interval (T8) (Elbau et al., 2018; Kühnel et al., 2020).

##### Neural response

To assess the average neural response to stress across the whole brain, we built a first-level model as previously reported (Kühnel et al., 2020). In this model, three regressors modeled the five arithmetic blocks (60s each) of the *PreStress*, *Stress* and *PostStress* condition, respectively. To account for motor responses, the model additionally included one regressor modeling individual motor responses. Moreover, the verbal feedback during the *Stress* phase was captured in one other regressor. To account for noise components, the models included the six movement parameters, their temporal derivatives, and the aCompCor components (physiological noise components (5 each) from white matter and cerebro-spinal fluid). Data were high-pass filtered with a cut-off of 256s. To assess the neural stress response and stress recovery we estimated the first level contrasts of interest, *Stress* – *PreStress*, and *PostStress* – *PreStress*, for each participant separately.

#### 2.4.3 Associations of stress responses with BMI

Across all stress responses (i.e., subjective, endocrine, autonomous, and neural), we used multiple regression (voxel-wise for neural stress responses) analyses to assess associations with BMI. All analyses additionally included age (associations with BMI, see Table S3), sex, pre-task cortisol, and diagnosis status (fulfilling the criteria for a F3 or F4 diagnosis within the last 12 months, (Kühnel et al., 2022)) as confounding variables. Moreover, we explored sex-specific associations of BMI with stress responses by including an interaction term. Analyses regarding the neural stress response (whole-brain regressions and predictive modeling), additionally included average log-transformed framewise displacement (Power et al., 2014) as a covariate, since a higher BMI has been associated with increased movement during scanning (Ekhtiari et al., 2019).

#### 2.4.4 Predictive modeling of BMI based on dynamic neural trajectories

To evaluate the predictive performance of stress-induced changes for the outcome BMI on an individual level, we used a recently published pipeline that captures dynamic trajectories of activation and functional connectivity changes between pre-defined regions of interest (ROI, (Kühnel et al., 2022)). Briefly, average timeseries (unsmoothed) were extracted from the preprocessed task and resting states in 21 ROIs (Shen et al., 2013) of a stress-related network including the left and right amygdala, hypothalamus, caudate, putamen, anterior, medial, and posterior hippocampus, anterior and posterior insula and one region for the posterior cingulate, dorsal anterior cingulate, and ventromedial prefrontal cortex. After denoising (detrending, despiking, and residualization with the same regressors as in the whole-brain analyses) and concatenation of the timeseries, hierarchical models for each edge (Kroemer et al., 2016; Kühnel et al., 2022; McLaren et al., 2012) were used to estimate block-wise changes in activation in 21 ROIs and their functional connectivity (FC, 21*20/2 edges). Each model then included the timeseries of one region (ROI_1_) as dependent variable and the timeseries of the other (ROI_2_) as independent variable. Moreover, they included separate regressors for each of the 15 task blocks and their interaction with the predicting timeseries to capture FC changes. Additionally, we again accounted for changes in activation corresponding to motor responses or verbal feedback, by including the two corresponding convolved regressors from the first-level models. Predictors for interaction terms were mean-centered. In all models, the predictors capturing changes in activation and FC were entered as random effects by participant, so that group-level and regularized individual-level estimates were calculated (Mejia et al., 2018; Narayan and Allen, 2016; Pervaiz et al., 2020). Next, individual estimates for changes in activation or FC were extracted from each model and aggregated either across regions (activation: combining bilateral regions) or four previously reported subnetworks showing similar stress response trajectories across the task (connectivity: (Kühnel et al., 2022)) leading to feature sets including between 68 (connectivity), 180 (activation) and 248 (activation + connectivity) features.

Last, we used those block-wise features to predict BMI with elastic net (*lasso*, preset alpha=.5, Matlab2020a) and nested 10-fold cross-validation. As in previous studies (Kühnel et al., 2022), we used elastic net since it performs well if the number of features is relatively high and they are correlated (Jollans et al., 2019). Confounding variables were included in baseline prediction models, and we evaluated the incremental variance explained by fMRI features. Statistical significance was determined using permutation tests (iterations=1,000; outcome was shuffled with confounders to keep their correlation).

#### 2.4.5 Contribution of peripheral inflammation

To determine the contribution of peripheral inflammation levels to changes in stress-induced brain responses associated with a higher BMI, we first selected cytokines that were correlated with BMI using partial correlations correcting for age, self-reported sex and presence of a psychiatric disorder. As we used this step only to select cytokines for further analysis, we did not correct for multiple testing but used an uncorrected p-value threshold of p < .05. Next, we evaluated whether each of the selected cytokines was associated with either the BMI predicted by the brain response or the residual BMI capturing variance not related to the brain response. Then, if a cytokine is correlated with the predicted BMI, this would indicate that both the stress-induced brain response and cytokine concentration explain shared variance in BMI. In contrast, if it is solely associated with the residual the cytokine and the stress-induced brain response explain independent variance in BMI. To this end, we used separate multiple regression models for each target cytokine predicting either the predicted BMI or the residual BMI, and age, presence of psychiatric diagnosis, pre-task cortisol response, sex, the cytokine and the interaction between sex and the cytokine as covariates.

#### 2.4.6 Statistical threshold and software

Statistical analyses were performed in Rv4.0.2 (R Core Team, 2018). For whole-brain fMRI analyses, the voxel threshold was set at *p_uncorrected_*<.001. Clusters were considered significant with a cluster-corrected threshold of *p_cluster.FWE_*<.05.

## 3. Results

### Higher stress-induced negative affect in high BMI participants

As reported in a previous publication (Kühnel et al., 2022), the task elicited a robust multi-level stress response (Figure 1) as indicated by an increase in heart rate (*b* = 6.7, *p* < .001), negative affect (*b* = 7.7, *p* < .001) and a decrease in positive affect (*b* = -2.2, *p* < .001). Cortisol increased in response to stress in participants not showing an increase in cortisol to the blood drawing procedure (*b* = 0.4, *p* = .011). After stress, heart rate recovered but not to baseline levels (*b* = 0.87, *p* = .034). Moreover, after a thirty-minute break, negative affect (*b* = -1.0, *p* = .014) but not positive affect (*b* = -1.3, *p* < .001) had recovered.

**Figure 1:**
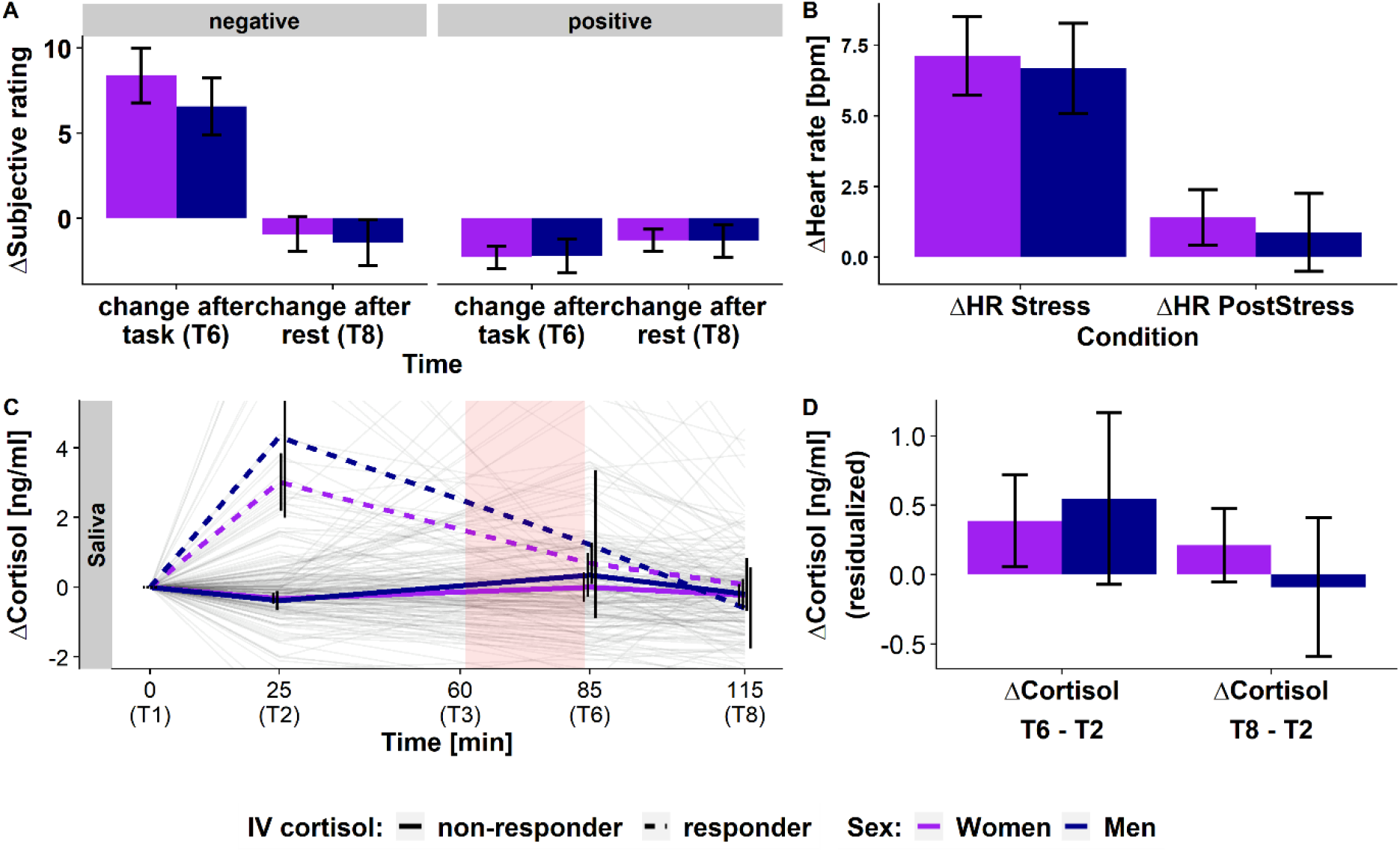
The stress task induced a similar stress response on the endocrine, subjective, and heart rate level in women and men. A) Stress-induced increases, relative to T2, (T6: *b*=7.7, *p*<.001) in negative affect that recover below baseline levels after stress (T8: *b* = -1.0, *p* = .014). At the same time, positive affect decreases (T6: *b*=-2.2, *p*<.001) but does not recover back to the levels at the beginning (T8: *b* = -1.3, *p* < .001). B) Stress induces increases in heart rate (*b* = 6.7, *p* < .001) that decreases during PostStress (*b* = -5.8, *p* < .001) but does not completely reach PreStress levels (*b* = 0.9, *p* = .033). C) Stress induces a cortisol response in participants not already reacting to the placement of an intravenous catheter (“non-responder”) compared to the pre-task cortisol measurement (T2). Cortisol levels recover close to baseline levels after the 30-minute break. Thin lines depict individual cortisol trajectories; thick lines show group averages. The shaded area shows the timing of the stress task. D) Changes in cortisol directly after the task (T6 – T2) and after the 30-minute rest (T8 – T2) do not differ between men and women. Values in A,B, and D show residualized (age, sex, IV-cortisol response, diagnosis status) averages and confidence intervals (95% CI) for men and women separately.

Next, we evaluated the effect of sex and BMI on stress reactivity. On the subjective level, a higher BMI was associated with higher stress-induced changes in negative affect directly after the task (*b* = 1.48, *p* = .047, N_women_= 120, Figure 2) and after the 30-minute rest period (*b* = 1.18, *p* = .019), relative to the baseline at T3. Notably, this association was significant in women at the second post stress time point (T6: *b* = 1.7, *p* = .069; T8: *b* = 1.3, *p* = .025) but not in men (T6: *b* = -1.1, *p* = .24; T8: *b* = 0.6, *p* = .44; Figure 2B). However, the interaction between sex and BMI did not reach significance (T6: *p* = .095; T8: *p* = .66). In contrast, higher BMI was not linked to stress-induced changes in positive affect, heart rate, or cortisol concentrations (*p*s > .10, Figure 2A, Table S4) and subjective, cardiovascular, and endocrine stress responses did not differ between men and women (*p*s > .15; Figure 2A, Table S2).

**Figure 2:**
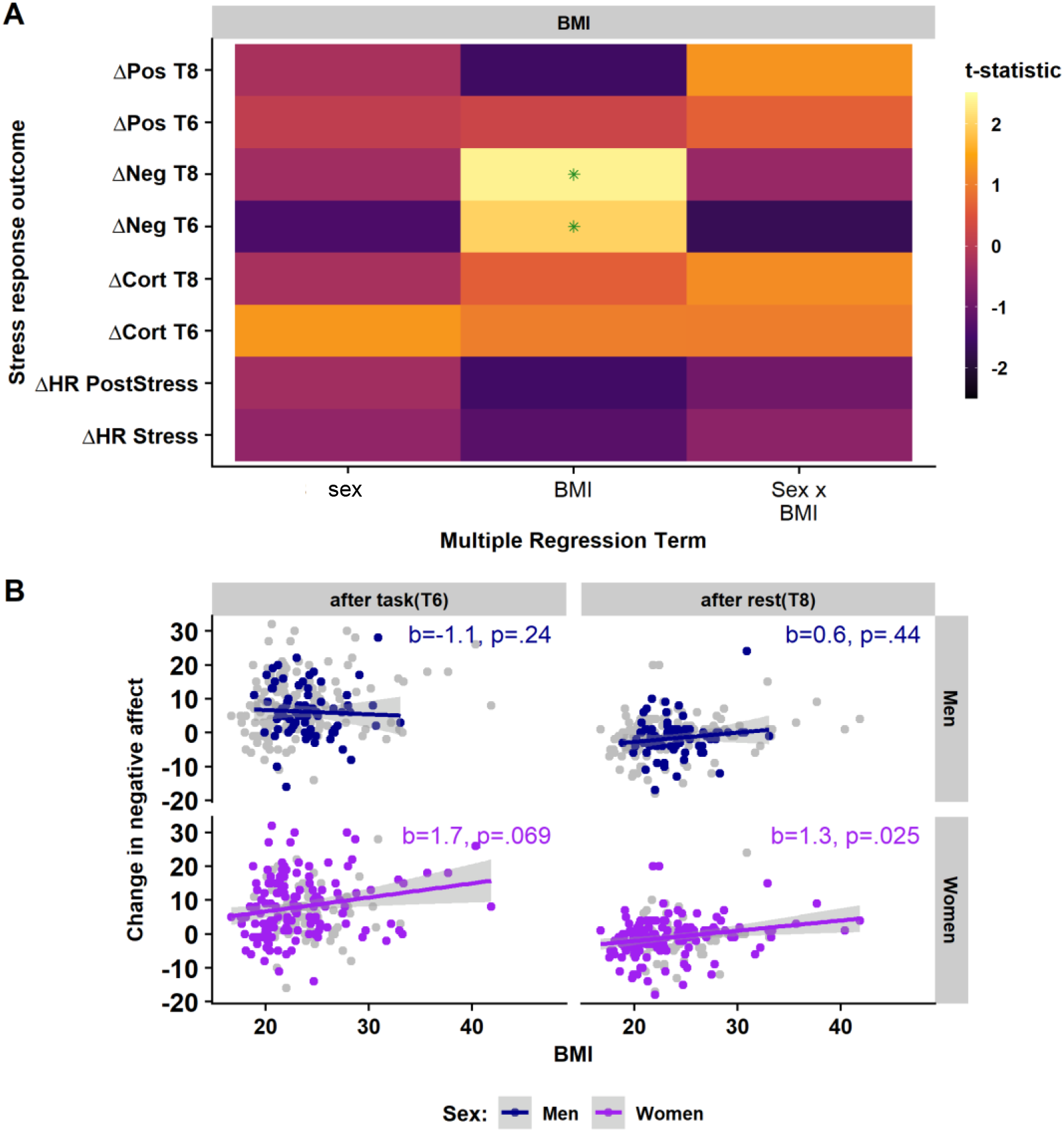
Stress-induced increases in negative affect are higher in participants with a high body mass index (BMI). A) Regression estimates for the effects of sex, BMI, and their interaction in models predicting multi-level stress responses. Estimates are t-values from linear multiple regressions adjusted for linear effects of age, presence of a psychiatric diagnosis, and cortisol response to intravenous catheter placement. Green asterisks indicate significant results (*p* < .05). B) Scatterplots showing the significant relationship between BMI and stress-induced negative affect after the task (both sexes: *b* = 1.48, *p* = .047) and after the following 30-minute rest (both sexes: *b* = 1.18, *p* = .019) separated for men and women to depict potential sex differences. Associations with BMI were significant in women (T8), but not in men. ΔCort T6, or T8 = Cortisol increase after the end of the task (T6) or after the additional rest period (T8), respectively, compared to baseline (T0). Again models include age, presence of a psychiatric diagnosis, and cortisol response to intravenous catheter placement as covariates but data is unadjusted for the visualization in the scatterplots. ΔHR PostStress = Difference in heart rate between task-block in the PostStress and PreStress condition, ΔHR Stress = Difference in heart rate between task-block in the PostStress and PreStress condition, ΔNeg T6 = Difference in state negative affect directly after the task (T6) compared to before the task (T3), ΔPos T6 = Difference in state positive affect directly after the task (T6) compared to before the task (T3). Same for T8, referring to the comparison with the time point after 30 minutes of rest.

### Reduced stress-induced activity in the substantia nigra and posterior insula in participants with high BMI

After establishing an association between BMI and stress-induced negative affect, we investigated whether the brain response to stress was also affected in participants with high BMI. As previously shown (Elbau et al., 2018; Kühnel et al., 2022, 2020), the social stress task led to robust stress-induced increases in BOLD responses in the visual and parietal cortex (i.e., task positive regions) as well as decreases in regions of the default mode network, including the posterior cingulate cortex (PCC) and angular gyrus. Moreover, stress reduced BOLD responses in the insula (posterior and anterior), supplementary motor area (SMA) and dorsomedial prefrontal cortex (dmPFC) activation.

In line with changes in negative affect, higher BMI was associated with stronger stress-induced decreases of BOLD responses in the posterior insula (L: *p*_FWE_ < .001, *k* = 293, R: *p*_FWE_ = .041, *k* = 143) and a cluster overlapping with substantia nigra (*p*_FWE_ < .001, *k* = 390) as well as increased BOLD responses in the precuneus/superior parietal lobe (*p*_FWE_ = .025, *k* = 145, Figure 3, maps have been uploaded to NeuroVault for inspection: https://neurovault.org/collections/NABGNECT/). To better characterize the associations of BMI and stress-induced activation, we extracted average beta values from ROIs (Shen et al., 2013) containing the significant clusters and performed post-hoc analyses. As for negative affect, the association with BMI was only significant in women (substantia nigra (SN): *b* = -0.05, *p* = .004, posterior insula R: *b* = -0.06 *p* < .001, posterior insula L: *b* = -0.03, p = .008) but not men (SN: *b* = -0.001, *p* = .95, posterior insula L: *b* = -0.03, *p* = .069, posterior insula R: *b* = 0.005, *p* = .75, Figure 3B). However, the interaction of sex and BMI did not reach significance (posterior insula R: *p* = .071, SN: *p* = .36), suggesting that the effect may be more pronounced in women, although a bigger sample is necessary to draw conclusions about the specificity of the effect for women.

**Figure 3:**
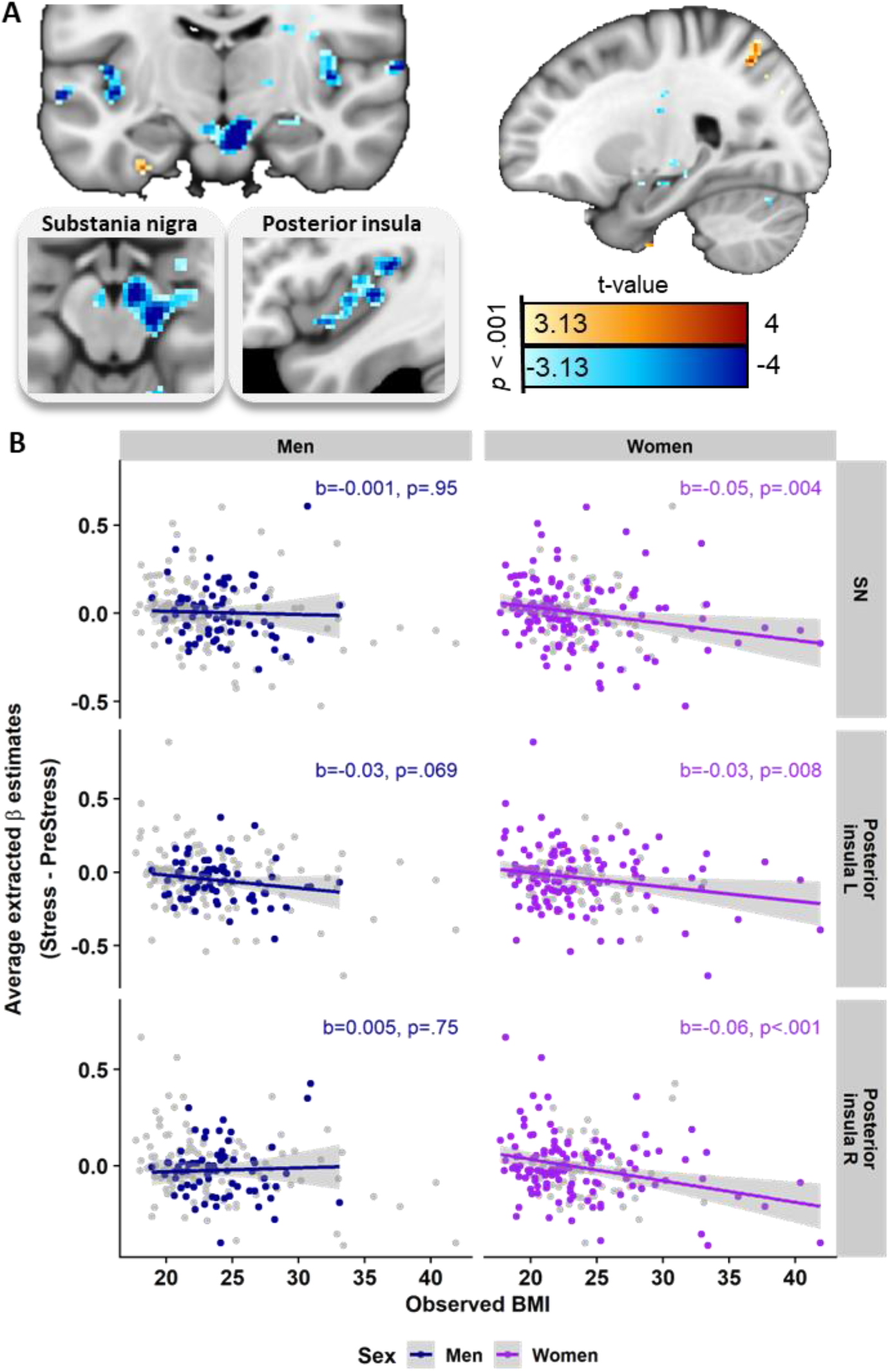
Stress-induced deactivations in the posterior insula and substantia nigra are stronger with increasing BMI. A) Whole-brain regression analyses showing associations between body mass index (BMI) and stress-induced (*Stress – PreStress*) activation changes. Higher BMI is associated with increased (warm colors) stress-induced activity in the superior parietal lobe / Precuneus and decreased (cool colors) activity in the substantia nigra and posterior insula. Voxel-threshold for display: p <.001, t > 3.13. B) Extracted beta estimates (average across the region of interest, ROI) from corresponding ROIs defined in the Shen Atlas (Shen et al., 2013) are negatively associated with the BMI. Regression weights and significance values are derived from separate multiple regressions for women and men also accounting for age, diagnosis of a psychiatric condition, and medication status but data is unadjusted for the visualization. The associations are only significant in women. BMI = body mass index, SN = substantia nigra, L = left, R = right.

### Stress-induced trajectories of brain activation predict BMI in women

As we previously showed that spatio-temporal profiles of stress-induced changes in brain activation and connectivity recover interindividual differences in stress-related phenotypes beyond conventional analyses (Kühnel et al., 2022) and because BMI has been associated with altered FC (Farruggia et al., 2020), we next tested whether those stress-induced spatio-temporal trajectories were also predictive of BMI on an individual level. To evaluate the predictive accuracy of individual response profiles, we used a recently established framework predicting BMI from stress-induced changes in FC, activation, or FC and activation combined (activation and connectivity changes across the 12 ROIs and 17 timepoints: 2 resting states before and after stress, 15 task blocks; FC of 4 clusters showing differential changes across task (Kühnel et al., 2022)). We predicted BMI using an elastic net algorithm with nested 10-fold cross-validation with an approach previously described in Kühnel et al. (2022). We predicted BMI based on activation (Δ*R*^2^ = .07, *p*_perm_ < .001, Figure 4A-E). Predictions based on FC changes or FC and activation changes combined did not predict BMI better than a baseline model including age, psychiatric diagnosis, and average framewise displacement (log-transformed). A higher BMI was predicted by higher activation of the anterior hippocampus, ventromedial prefrontal cortex, and dorsal anterior cingulate cortex (dACC) as well as lower activation of the posterior insula and posterior hippocampus mirroring the whole brain-associations. In line with the conventional analyses reported above, the prediction of the elastic net was only better than chance in women (women: *r* = .26, *p* = .005; men: *r* = -.05, *p* = .66, Figure 4A). Re-training and evaluating the model only in women further numerically improved the predictive value (Δ*R*^2^ = .11; *p*_perm_ = .002, compared against a baseline model including confounding variables, Figure 4F), although selected features were similar across both models (Figure 4C, SI) with slightly higher weights in the retrained model including only women.

**Figure 4:**
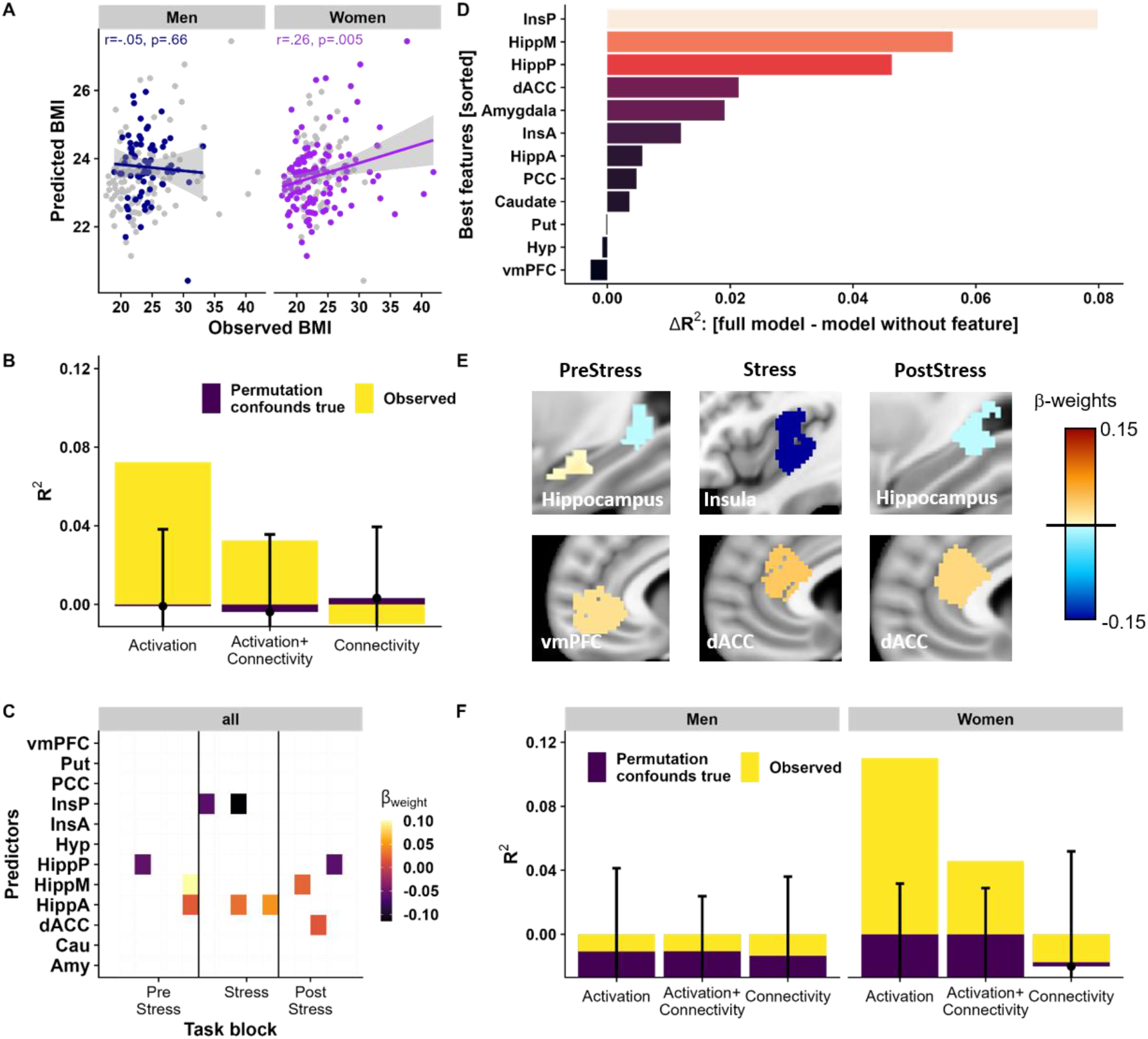
Block-wise changes in activation across the task predict body mass index (BMI) in women. A) An elastic net model based on activation changes across task-blocks predicts BMI. Predicted and observed values of BMI were significantly correlated across the complete sample (*r*=.33, *p*perm<.001). This association was driven by women (*r*=.26, *p* = .005), but was not seen in men (*r*= -.05, *p* =.66). Prediction models included covariates (age, sex, diagnosis, pre-task cortisol, and log-transformed average framewise displacement), but for visualization, data is unadjusted. B) The model based on stress-induced activation trajectories (yellow) predicted BMI beyond a baseline model based on confounding variables (age, diagnosis, pre-task cortisol, log-transformed average framewise displacement, and medication). The observed R2 (yellow) is higher than the 95% percentiles of the model with true confounds but permuted features (repeated 1,000 times). In contrast, models based on functional connectivity (FC trajectories), or a combination of FC and activation trajectories did not perform better than the baseline model including confounds (i.e., observed R2 within 95% percentile range). Overlapping bars show the average model performance of the observed model (yellow) or the baseline models with permuted features (violet). Error bars depict 95% percentiles derived from permuting the outcome together with the confounding variables to evaluate the contribution of the activation features beyond the confounds. Av C) Standardized weights from the prediction model including stress-induced changes in activation. Depicted weights were retained in ≥80% of outer cross-validation folds. D) Importance of each feature set (i.e., all timepoints of one region) for the prediction of BMI. The *ΔR^2^* reflects how much predictive accuracy is lost when leaving out all timepoints of the feature E) Standardized weights predicting BMI across the complete sample in the model including averaged activations for *PreStress*, *Stress*, and *PostStress*. F) Activation changes (and changes in activation and FC combined) only predict BMI in women, but not men, beyond a baseline model including confounds (age, diagnosis, pre-task cortisol, log-transformed average framewise displacement, psychopharmacological treatment) when training separate models. Error bars depict 95% percentiles derived from permuting the outcome together with confounds to evaluate the contribution of features beyond confounds. vmPFC = ventromedial prefrontal cortex, Put = Putamen, PCC = posterior cingulate cortex, InsP = posterior insula, InsA = anterior insula, Hyp = Hypothalamus, HippP = posterior hippocampus, HippM = medial hippocampus, HippA = anterior hippocampus, dACC = dorsal anterior cingulate cortex, Cau = caudate, Amy = amygdala

### Baseline cortisol, but not cytokines, are associated with predicted BMI

Last, we evaluated whether increased inflammation in overweight and obesity might explain the observed alterations in stress reactivity. To this end, we first identified cytokines that are associated with higher BMI using partial correlation analyses correcting for age, diagnosis status, and current psychiatric medication. In line with previous studies (Benson et al., 2009; Caslin et al., 2016; Mathieu et al., 2010), higher BMI was associated with increased high sensitivity C-reactive protein (hsCRP), interleukin (IL)-1 receptor antagonist (IL-1RA), and vascular endothelial growth factor A (VEGF-A), intracellular adhesion molecule 1 (ICAM-1), chemokine (C-C motif) ligand 2 (MCP-1), chemokine (C-C motif) ligand 4 (MIP-1beta), chemokine (C-C motif) ligand 22 (MDC), tumor necrosis factor (TNF-alpha), IL-16, serum amyloid A (SAA), and soluble IL-6 receptor (sIL-6R, Figure S2). Moreover, baseline cortisol (measured in plasma samples at the beginning of the first session ∼8 am) was lower in participants with high BMI (rho = -.17, p = .015). Again, this was driven by a stronger association in women (rho = -.27, p = .003 vs. men: rho = -.15, p = .22), but associations did not differ significantly between sexes (interaction; *p* = .64). Of note, baseline plasma cortisol measured on a separate day as the stress task was correlated (r = .37, p <.001, Figure S3) with the first salivary cortisol measurement on before the stress task, but after other components of the session.

To determine whether stress-induced changes in brain function that were predictive of BMI were also linked to BMI-related increases in inflammation, we evaluated if cytokines were correlated with predicted versus residual BMI. The rationale for this analysis is that associations of cytokines with predicted BMI would indicate shared variance, whereas associations with residual BMI would indicate that variance attributable to cytokines is not captured by the predictive model based on brain function. In fact, cytokines were only associated with residual BMI, whereas reduced baseline cortisol levels were also associated with a higher predicted BMI (BMI: *b* = -0.31, *p* = .023; BMI x Sex: *p* = .086, Figure 5), suggesting that obesity-related peripheral inflammation does not account for the BMI variance predicted by stress-induced brain responses.

**Figure 5:**
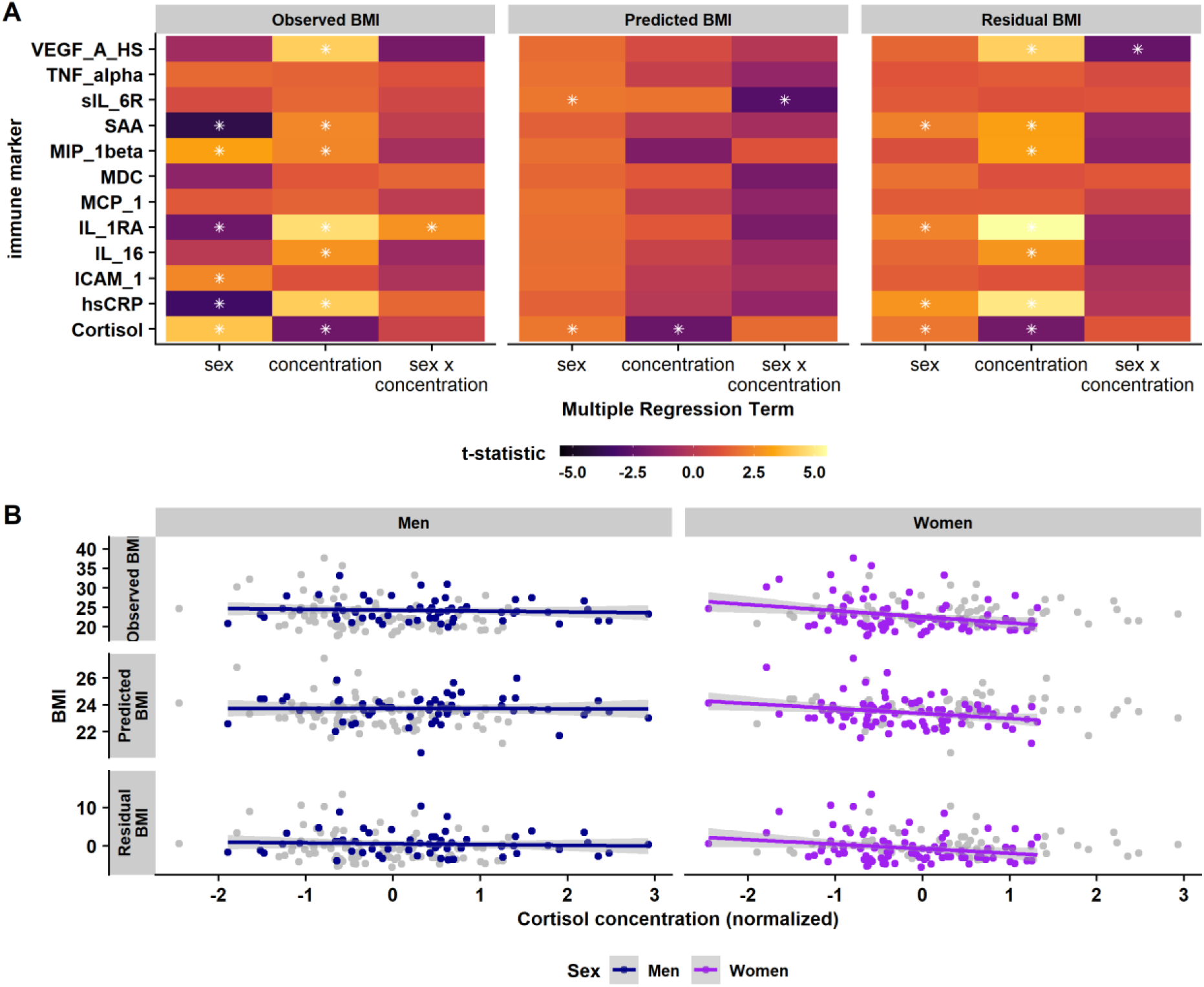
Peripheral levels of cytokines do not account for associations with predicted BMI based on stress-induced brain response patterns. A) Multiple regression coefficients for the associations between sex, normalized cytokine concentration, and their interaction with participant’s actual BMI, the BMI predicted by stress-induced activation trajectories, and the residual BMI. Only baseline (morning) cortisol concentration was related to the observed as well as predicted BMI. White asterisks indicate significant predictors. All regressions models include age, psychiatric diagnosis, and medication status as additional covariates B) Scatterplots for the association between baseline cortisol and BMI (observed, predicted, and residual), separated for sex. Associations were only apparent in women (rho = -.27, p = .003), not men (rho = -.15, p = .22). However, the interaction between sex and cortisol concentration was not significant (*p* = .64). Correlation values are partial correlations corrected for age, diagnosis status and current psychiatric medication, but data is unadjusted for the visualization in the scatterplots. hsCRP = high sensitivity CRP, IL-1RA = interleukin 1 receptor antagonist, VEGF-A = vascular endothelial growth factor A, ICAM-1 = intracellular adhesion molecule 1, MCP-1 = chemokine (C-C motif) ligand 2, MIP-1beta = chemokine (C-C motif) ligand 4, MDC = chemokine (C-C motif) ligand 22, TNF-alpha = tumor necrosis factor alpha, IL-16 = interleukin 16, SAA = serum amyloid A, and sIL-6R = soluble IL-6 receptor.

## 4. Discussion

Stress has been related to an increased risk for overweight and obesity, and in turn a higher BMI is associated with changes in acute stress reactivity. However, the contribution of increased peripheral inflammation and changes in the HPA axis are not yet well understood. Here, we demonstrate that participants with a higher BMI show increased stress-induced changes in negative affect accompanied by a stronger stress-induced brain response in the posterior insula, substantia nigra, and parietal cortex. Accordingly, BMI was predicted by activation trajectories across stress phases derived from the posterior insula, the dACC, and the hippocampus (anterior and posterior). This model was only predictive in women and associations of acute stress with BMI were primarily driven by variance in women. Crucially, different levels of cytokines did not account for altered stress reactivity in the brain as they were uncorrelated with predicted BMI, whereas baseline cortisol was associated with the predicted BMI based on brain responses to stress. To summarize, our results provide initial evidence that acute stress reactivity is more strongly interrelated with BMI in women compared to men and that these altered responses are more strongly linked to endocrine changes compared to cytokine changes associated with overweight and obesity.

A higher BMI was predicted in the whole sample by lower stress-induced activation in the posterior insula and posterior hippocampus as well as higher activation in the anterior hippocampus and dACC. Both the hippocampus (Elbau et al., 2018; Herman et al., 2005; Pruessner et al., 2008) and the dACC (Eisenberger et al., 2007; McEwen and Gianaros, 2010; Sandner et al., 2020) have been repeatedly associated with stress and emotion regulation. On the one hand, increased activation of the hippocampus during stress with a higher BMI suggests altered regulation of the HPA axis as brain responses of the hippocampus have been associated with stress-induced cortisol responses (Elbau et al., 2018; Pruessner et al., 2008). Likewise, hippocampal activation while viewing aversive pictures was negatively associated with a baseline cortisol index (Cunningham-Bussel et al., 2009). Such an interpretation is strengthened by lower baseline cortisol in participants with high BMI as baseline cortisol levels in turn affect stress-induced cortisol responses (Gossett et al., 2018; Kudielka et al., 2004). However, stress-induced cortisol responses were unrelated to BMI in our sample in line with previous mixed evidence (Herhaus and Petrowski, 2018; McInnis et al., 2014; Therrien et al., 2010). Increased activation of the dACC with a higher BMI might be related to an increased recruitment of the salience network in response to stress (van Oort et al., 2017). Notably, dACC activation was predictive of BMI during Stress and PostStress, suggesting a broader role of the dACC in performance monitoring in participants with high BMI. On the other hand, lower activation of the posterior insula during stress with a higher BMI might be related to reduced integration of interoceptive signals and a higher focus on exteroception (Barrett and Simmons, 2015; Craig, 2009; DeVille et al., 2018; Farb et al., 2013; Kuehn et al., 2016; Livneh et al., 2020). Comparably, the posterior insula has been implicated in the processing of negative social feedback (Miedl et al., 2016) differentially for men and women (Lee et al., 2014). Importantly, ROI-based results from the predictive model were in line with conventional whole-brain analyses, where higher BMI was related to stronger stress-induced decreases of activation in the posterior insula. Moreover, we previously showed that posterior insula activity was also predictive of negative affectivity (Kühnel et al., 2022), a trait related to anxiety and depression, which, as obesity, are also related with chronic stress and have a shared genetic basis with obesity as well changes in the immune system (Kappelmann et al., 2021). To conclude, our results point to an exaggerated stress response with higher BMI in regions implicated in interoceptive and exteroceptive (salience) processing and related to regulation of the HPA axis.

Differences in stress-induced brain responses with high BMI might be related to changes in the endocrine as well as the immune system and their interactions (Incollingo Rodriguez et al., 2015; Mathieu et al., 2010). Stress-induced cortisol responses were not related to BMI, contrasting earlier studies (Benson et al., 2009; Dockray et al., 2009; Herbison et al., 2016), although recent evidence has been inconclusive (Herhaus and Petrowski, 2018; McInnis et al., 2014; Therrien et al., 2010). One possible explanation is that although highly correlated, not BMI but visceral adiposity is differentially related to HPA-axis functioning (Incollingo Rodriguez et al., 2015). Instead, in line with previous studies (Champaneri et al., 2013; Schorr et al., 2015), high BMI was associated with lower baseline cortisol levels, especially in women. Critically, lower baseline cortisol levels were also associated with predicted BMIs based on the trained elastic net model pointing to shared variance with stress-induced changes in activation. This fits well with other studies on for example hippocampal responses to stress (Elbau et al., 2018; Pruessner et al., 2008) since baseline cortisol level might influence stress reactivity (Chen et al., 2017; Juster et al., 2012; Kudielka et al., 2004; Kühnel et al., 2020). Likewise, exogenous administration of corticosteroids has been associated with changes in a network associated with the regulation of ingestive behavior, which includes the hippocampus, insula, hypothalamus, and related cortico-striatal regions (Bini et al., 2022; Symonds et al., 2012). Taken together, our results point to an altered set point of the HPA axis in obesity that is also related to altered neural stress reactivity.

In line with previous studies (Caslin et al., 2016; Mathieu et al., 2010), participants with a higher BMI showed higher levels of peripheral inflammation, but the predicted BMI based on stress-induced activation trajectories was independent of changes in peripheral cytokines. This indicates that the influence of higher obesity-related inflammation on acute stress reactivity (Buzgoova et al., 2020; Chiappelli et al., 2016; Kunz-Ebrecht et al., 2003) is independent and potentially additive to effects on other systems, such as the HPA axis. To summarize, our results suggest that modeling of stress-induced changes in the brain can also help to unravel potential obesity-associated alterations across various systems such as the immune or stress response system.

Of note, prediction of BMI based on stress-induced activation trajectories was only successful in women. Accordingly, women also showed stronger associations between BMI and activation in the posterior insula, the subjective stress experience, and baseline cortisol. Overweight and obesity are more prevalent in women (Garawi et al., 2014; Hedley et al., 2004) and women show distinct stress responses (Kuhn et al., 2023). Moreover, increased food intake in response to stress is more often reported by women (Adam and Epel, 2007; Grunberg and Straub, 1992; Meule et al., 2018; Zellner et al., 2006). Notably, negative affective responses to the stressor are related to compensatory increased food intake (Macht and Mueller, 2007) in the same way that altered endocrine stress reactivity (Herhaus et al., 2020; Newman et al., 2007) is also thought to affect food intake. Therefore, our results emphasize the role of acute and chronic stress in obesity and overeating.

Despite the notable strengths of our study, it has several limitations that call for additional research. First, the sample size is comparatively large for a task-based neuroimaging study, but slightly imbalanced with two thirds of the sample being women, roughly reflecting the different incidence of mood and anxiety disorders. The sample covers a broad range of BMI (18 – 41 kg²/m) in women, while the range is more restricted in men (20 – 33 kg^2^/m). Since negative effects of overweight and obesity on the immune system or energy metabolism are conceivably larger in obesity (i.e., BMI > 30 kg/m², Monteiro and Azevedo, 2010; Wisse, 2004; Yudkin, 2007), a replication of sex-specific effects in a larger and more balanced sample is necessary. Likewise, associations with peripheral markers of inflammation might only be apparent in samples with more participants with obesity. Second, we only assessed peripheral inflammation markers, although central inflammation might have distinct mechanisms affecting BMI and stress reactivity (Chan et al., 2019). Third, we did not account for effects of the menstrual cycle or the use of hormonal contraception in women, although stress reactivity has been shown to be affected by the hormonal state (Childs et al., 2010; Kirschbaum et al., 1999). Moreover, other lifestyle factors such as smoking or alcohol consumption might affect associations between BMI and cytokine levels. Fourth, likewise while sessions all started at the same time to account for circadian effects on stress reactivity, we did not standardize the metabolic state of participants, although the current glucose levels have been shown to affect stress reactivity (von Dawans et al., 2021) and energy metabolism might differ across the BMI. Fifth, our results are cross-sectional so we cannot draw conclusions about causal relationships between stress reactivity, overweight and obesity, and associated changes in endocrine or immune systems. Sixth, we only investigated associations with self-reported sex, which in our sample was congruent with the sex determined by genotyping. However, effects based on gender in samples including more diverse populations might differ.

To summarize, we show that BMI can be predicted by stress-induced activation trajectories of the insula, hippocampus, and dACC in women but not in men. In line with predictive modeling of brain responses, participants with high BMI showed stronger stress-induced changes in negative affect and lower baseline cortisol levels. This may indicate a changed set point of the HPA axis in participants with high BMI, which might contribute to altered stress reactivity and might point to shared alterations with other stress-related disorders such as depression that also have overlapping genetic factors (Kappelmann et al., 2021). To conclude, our results point to an important role of the HPA axis and acute stress reactivity in the mechanisms underlying obesity and overweight, particularly in women. If validated, such sex differences in altered stress reactivity with a higher BMI may contribute to sex differences in obesity as well as stress-related eating more broadly.

## Acknowledgement

We thank Anna Hetzel and Ines Eidner for help with data acquisition, Manfred Uhr and the team of the Max Planck Institute of Psychiatry Biobank for sample processing. The BeCome working group including Tanja Brückl^1^, Victor I. Spoormaker^1^, Angelika Erhardt^1^, Norma C. Grandi^1^, Julius Ziebula^2^, Immanuel G. Elbau^1^, and Susanne Lucae^2^ were responsible for conception, organization, and execution of the study. Nils B. Kroemer received support from the Deutsche Forschungsgemeinschaft (DFG) grants KR 4555/7-1 and KR 4555/9-1. Victor I. Spoormaker has received income from consultations and advisory services for Roche. Knauer-Arloth was supported by the Brain Behaviour Research Foundation (NARSAD Young Investigator Grant, #28063).

^1^ Department of Translational Research in Psychiatry, Max Planck Institute of Psychiatry, Munich, Germany

^2^ Max Planck Institute of Psychiatry, Munich, Germany

## Author contributions

EBB, PGS and the BeCome working group were responsible for the concept and design of the BeCOME study. MC and PGS validated the paradigm and procedure. AK and NBK conceived the method and analysis plan and AK performed the data analysis. AK wrote the first draft of the manuscript and NBK contributed to the writing. JH, MK and JKA contributed the cytokine measures. All authors contributed to the interpretation of findings, provided critical revision of the manuscript for important intellectual content and approved the final version for publication.

