## Supplementary Information for "Stress-induced brain responses are associated with BMI in women"

##### Assessment of subjective stress experience (BSKE scales):

The BSKE (*Befindlichkeitsskalierung durch Kategorien und Eigenschaftswörter*, Janke, 1994) scales are a short version of the more extensive "*Eigenschaftswörterliste*" (EWL, (Janke and Debus, 1978), a scale developed to assess current emotional state across positive and negative dimensions. This reduced scale consists of 15 items (emotions / states) relevant for anxiety that have been previously used to assess stress reactivity (Elbau et al., 2018; Ising et al., 2008; Kühnel et al., 2020) and comparable to the PANAS or state anxiety questionnaire that are also used in stress research (Kuhn et al., 2021; Shilton et al., 2017) assesses different emotions and feelings that might be affects by stress such as agitation, anxiety, anger, or sensitivity. Participants were asked to rate their current state/feeling ("I feel …") on 6-point scale ranging from 1 ("not at all / gar nicht") to 6 ("very strongly / sehr stark"). We calculated sum scores including the items activity, wakefulness, self-certainty, focus, and relaxed state of mind for positive affect and including the items internal and external agitation, anxiety, sadness, anger, dysphoria, sensitivity as well as three items assessing somatic changes for negative affect.

##### Concatenation of resting-state and task timeseries:

To assess task-induced functional connectivity changes referenced to a resting-state baseline we concatenated timeseries data from the psycho-social stress task and the two, flanking resting-states. Timeseries were linearly detrended (we did not include a quadratic trend to prevent excluding potential task effects with the same pattern induced by the task structure with non-stress phases flanking the acute stress), despiked, and denoised for each measurement separately so that the average gray scale values of each measurement was 0. To concatenate the task timeseries with the resting states, we matched the average gray scale values of the flanking resting-states with the average gray scale value of the rest baseline phases (fixation cross) during the *PreStress* condition for each region of interest. To this end, we calculated the average gray scale value (i.e., the measured raw intensity of the fMRI images) after detrending and denoising for each region of interest of the rest baseline phases (fixation cross) during the *PreStress* condition and then subtracted this offset from the complete task timeseries, so that the average intensity value during the rest baseline phases during *PreStress* was 0 and matched the average intensity values of the flanking resting-states.

##### Feature extraction for activation changes across task blocks:

To quantify changes in activation across task blocks, we extracted block-wise estimates from the same linear mixed-effects models we used for the dynamic changes in functional connectivity. Crucially, these models include regressors capturing task-induced changes in activation elicited by task structure (i.e., one regressor for each task block, one regressor for the motor response, and one regressor for the verbal feedback). Since we only estimated the upper triangle of the connectivity matrix, each region of interest was the target region in a different number of models (ranging from 20 to 1). For prediction we used an average across the 210 models based on their anatomical region and combined across all models predicting the same region and subsequently across left and right ROIs leading to 12 predictors (vmPFC, dACC, PCC, aIns, pIns, Caudate, Putamen, Amygdala, Hypothalamus, aHipp, mHipp, pHipp) for the stress phase prediction and 12x15 predictors (trajectories across time for all regions) for the prediction of interindividual differences.

### Figures

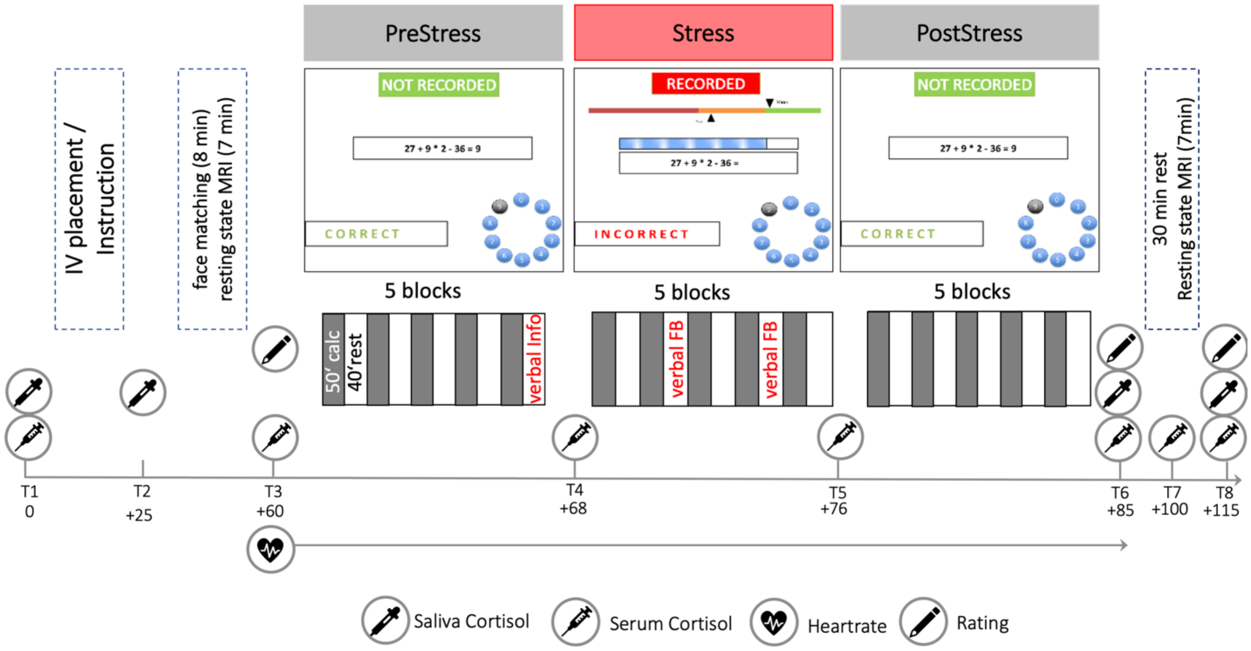


**Figure S1:** Detailed description of the psychosocial stress task (Kühnel et al., 2020). Before the stress phase, participants were informed about being recorded in the following trials. Additional aversive verbal feedback (verbal FB) about unsatisfactory performance was given in the 2nd and 4th rest period of the *Stress* condition. Saliva sampling was done in all participants (N=192) and in subsample of n=73 participants blood samples were taken to assess the cortisol response with higher temporal resolution.

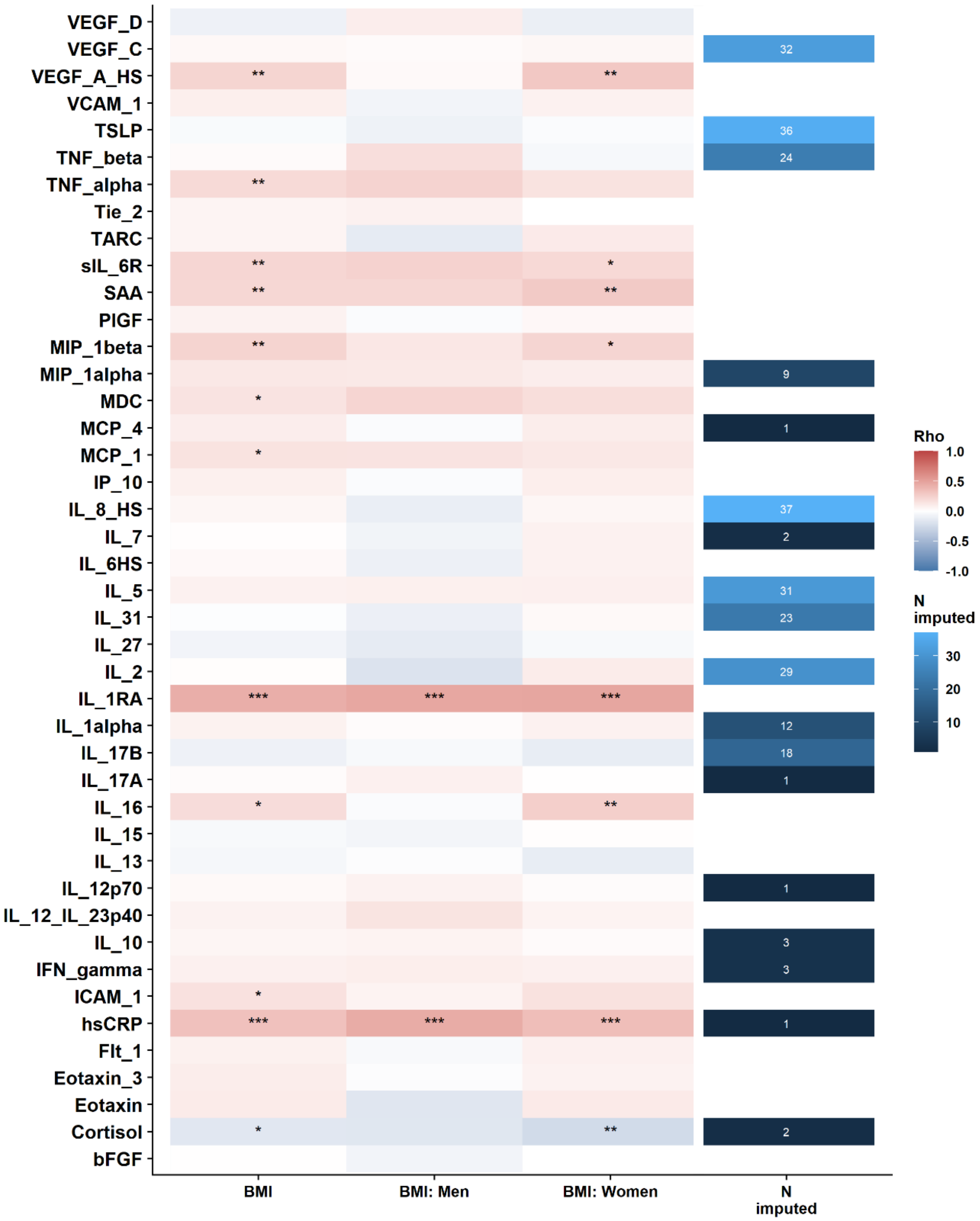


**Figure S2:** Uncorrected, partial correlations of body mass index (BMI) with all different immune markers, for the complete sample and men and women separately in a larger sample (N=198). All correlations are corrected for age, diagnosis, and current psychiatric medication. The last column shows the number of values that have been imputed. White indicates no values had to be imputed.

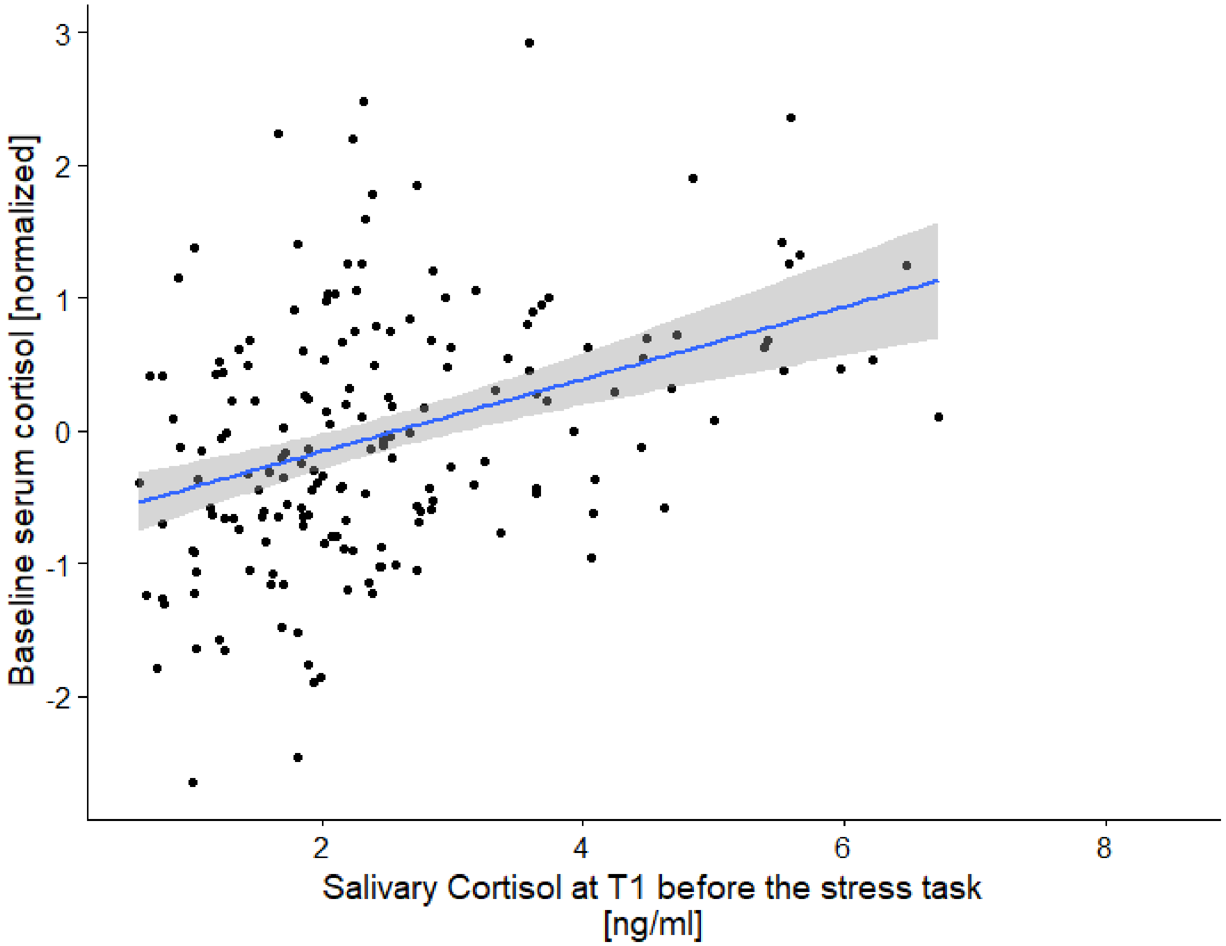


**Figure S3:** Correlation ($r$ = .38, $p$ < .001) of the baseline morning cortisol assessment (measured from a serum sample together with the cytokines) at separate day and the first salivary cortisol sample before the stress task. This sample was taken at approximately 10am and after participants had already completed a fear extinction paradigm.

### Tables

**Table S1**: Current and lifetime prevalence of psychiatric disorders identified using the CIDI in the present sample.

| | **12-months diagnosis N(%)** | **Lifetime diagnosis N(%)** |
|---|---|---|
| Substance use disorders (F1) | 8 (4%) | 42 (22%) |
| Mood disorders (F3) | 65 (35%) | 75 (39%) |
| Anxiety-related disorders (F4) | 101 (53%) | 122 (64%) |
| Other disorders | 8 (4%) | 14 (7%) |
| No diagnoses | 74 (39%) | 50 (26%) |
| 1 diagnosis | 56 (29%) | 54 (28%) |
| 2 diagnoses | 51 (27%) | 59 (31%) |
| 3 and more diagnoses | 9 (5%) | 27 (14%) |

*Note:* Anxiety disorders (F4) include specific phobias. Stress-related disorders in the last 12 months are defined as participants with a mood or anxiety-related disorder within the last 12 months excluding specific phobias. The control includes all other participants that might still receive diagnoses from other axes (e.g. substance use disorder: smoking).

**Table S2**: Number of participants included across all analyses.

| ***N*** | Subjective | Endocrine | Heart rate | Neural | Immune markers |
|---|---|---|---|---|---|
| BMI | 189 | 186 | 165 | 190 | 148 |

Note: Exclusion reasons: Endocrine (salivary cortisol) not enough material, Heart rate insufficient data quality

**Table S3**: No sex-specific effects of age on BMI

| Dependend variable | Predictors | Estimate | Std. error | p-value |
|---|---|---|---|---|
| BMI | Sex | -0.22 | 0.29 | 0.44 |
| | Age | 0.64 | 0.29 | 0.028 |
| | BMI * Age | -0.04 | 0.30 | 0.89 |

**Table S4**: Multiple regression models predicting stress responses (subjective, cardiovascular, endocrine) by sex, BMI, and their interaction.

| Group | Characteristic | Beta | 95% CI[1] | p-value |
|---|---|---|---|---|
| ΔHR PostStress | Sex | -0.39 | -2.0, 1.2 | 0.64 |
| | BMI | -0.61 | -1.6, 0.35 | 0.21 |
| | BMI * Sex | -0.66 | -2.9, 1.5 | 0.56 |
| ΔHR Stress | Sex | -0.38 | -2.5, 1.8 | 0.73 |
| | BMI | -1.0 | -2.2, 0.28 | 0.13 |
| | BMI * Sex | -1.4 | -4.3, 1.5 | 0.35 |
| ΔCortisol T6 | BMI | 0.19 | -0.17, 0.55 | 0.30 |
| | Sex | 0.37 | -0.25, 1.0 | 0.24 |
| | BMI * Sex | 0.44 | -0.42, 1.3 | 0.32 |
| ΔCortisol T8 | BMI | 0.09 | -0.16, 0.33 | 0.49 |
| | Sex | -0.10 | -0.51, 0.32 | 0.65 |
| | BMI * Sex | 0.35 | -0.23, 0.93 | 0.24 |
| ΔNegative affect T6 | **BMI** | **1.5** | **0.01, 2.9** | **0.047** |
| | Sex | -1.8 | -4.3, 0.66 | 0.15 |
| | BMI * Sex | -2.9 | -6.3, 0.55 | 0.10 |
| ΔNegative affect T8 | **BMI** | **1.2** | **0.20, 2.2** | **0.019** |
| | Sex: Men | -0.50 | -2.2, 1.2 | 0.56 |
| | BMI * Sex | -0.50 | -2.9, 1.9 | 0.67 |
| ΔPositive affect T6 | BMI | 0.09 | -0.60, 0.78 | 0.79 |
| | Sex | 0.23 | -0.94, 1.4 | 0.70 |
| | BMI * Sex | 0.52 | -1.1, 2.2 | 0.54 |
| ΔPositive affect T8 | BMI | -0.52 | -1.2, 0.15 | 0.12 |
| | Sex | 0.01 | -1.1, 1.1 | 0.98 |
| | BMI * Sex | 1.0 | -0.58, 2.6 | 0.21 |

[1]CI = Confidence Interval, BMI = Body mass index, In the multiple regression sex was dummy-coded with 0 = women and 1 = men. All models additionally included age, diagnosis status, cortisol response to the placement of an IV (responder = 1, non-responder 0) and medication status.
